# Adapting the Eysenck Personality Questionnaire-Revised Neuroticism scale for use in epidemiologic studies: A psychometric evaluation using item response theory in the UK Biobank

**DOI:** 10.1101/741249

**Authors:** Sarah Bauermeister, Chris Patrick Pflanz, John Gallacher, Dementias Platform UK

**Affiliations:** Department of Psychiatry, Medical Sciences Division, Warneford Hospital, University of Oxford, Oxford, United Kingdom

## Abstract

Neuroticism has been described as a broad and pervasive personality dimension or ‘heterogeneous’ trait measuring components of mood instability, such as worry; anxiety; irritability; moodiness; self-consciousness and sadness. Consistent with depression and anxiety-related disorders, increased neuroticism places an individual vulnerable for other unipolar and bipolar mood disorders and therefore highly relevant in epidemiologic research. However, the measurement of neuroticism remains a challenge. We aimed to adapt the 12-item Eysenck Personality Questionnaire-Revised Neuroticism (EPQ-RN) scale for use in epidemiologic studies by identifying psychometrically efficient items using item response theory. The 12-item EPQ-RN scale was evaluated by estimating an IRT model on data from 401,527 UK Biobank participants aged 39 to 73 years (*M* = 56.41 years; *SD* = 8.06), 53.68% female. The IRT model yielded two item characteristics: item discrimination, an indicator of how well an item differentiates between respondents, and item difficulty, an indicator of the amount of the latent construct (neuroticism) needed to endorse an item. The EPQ-RN exhibited psychometric inefficiency with poor discrimination at extremes of the scale range. High and low scores are relatively poorly represented and uninformative suggesting that high neuroticism scores derived from the scale are a function of cumulative mid-range values. Following systematic item deletion, a 3-item scale was found to have high levels of discrimination, but offered a narrow range of difficulty i.e. was not sensitive to low levels of neuroticism. A 7-item scale was found to be most informative; providing high levels of discrimination across the range of neuroticism scores.

## Introduction

Neuroticism has been operationally defined as a personality trait assessed by items referencing to instances of worry; anxiety; irritability; moodiness; self-consciousness; and sadness [1–3]. The NEO-PI (Neuroticism-Extraversion-Openness Personality Inventory) operationalises neuroticism as a combination of individual behavioural traits which may also be measured as isolated components of mood state e.g., anxiety; hostility; depression; self-consciousness; impulsiveness and vulnerability [4]. Eysenck has further argued that neuroticism is a direct reaction to the autonomic nervous system [5, 6], findings supported increased neuroticism correlated with tolerance to a highly stressed environment, suggesting a habituation relationship with everyday stressors [7, 8].

Eysenck’s attempts to define neuroticism and evaluate the measurement items thereof resulted in an original version of the Eysenck neuroticism scale existing as a component of the Maudsley Medical Questionnaire [9]. Assessment outcomes of this scale were reported in the Manual for the Maudsley Personality Inventory (MPI) [10]. Revision of the MPI by removing several items, and by using clinical judgement and factor analysis, has resulted in a revised neuroticism scale as a component of the Eysenck Personality Questionnaire [11], which has become a gold standard for neuroticism assessment and is therefore widely used in epidemiological research and cohort studies.

Using correlational techniques commonly used in classical test theory (CTT) for item deletion, has a bias towards identifying closely associated items as being informative. However, it is relatively opaque to the informativeness of individual items. The EPQ-R neuroticism scale (EPQ-RN), for example, has been found to lack items identifying respondents who would normally endorse items at the extreme ends of the trait continuum, e.g. high vs. low neuroticism [12].

We investigated the psychometric efficiency of the 12-item EPQ-RN [11] as a widely used measurement of neuroticism. We applied item response theory (IRT) to psychometrically evaluate the EPQ-RN using data from UK Biobank [13], a large population study which assessed neuroticism at baseline. Our expectation was that the large sample size and its heterogeneous population base would provide valuable item-level information for assessing the informativeness of individual items and the overall psychometric reliability of the scale.

## Methods

### Design and sample

We conducted a cross-sectional analysis of all UK Biobank participants providing EPQ-RN data during the baseline assessment. UK Biobank is a large population-based prospective cohort study of >500k participants [13]. Further details on design and procedure, including ethical approval, have been previously reported by Sudlow et al. [13].

### Assessment

The selection of mental health assessments was completed on a touchscreen computer, including the 12-item EPQ-RN [11] where participants were required to answer, ‘yes’, ‘no’, ‘I don’t know’ or ‘I do not wish to answer’ in response to the 12 questions: ‘Does your mood often go up and down?’; ‘Do you ever feel just miserable for no reason?’; ‘Are you an irritable person?’; ‘Are your feelings easily hurt?’; ‘Do you often feel fed-up?’; ‘Would you call yourself a nervous person?’; ‘Are you a worrier?’; ‘Would you call yourself tense or highly strung?’; ‘Do you worry too long after an embarrassing experience?’; ‘Do you suffer from nerves?’; ‘Do you often feel lonely?’; ‘Are you often troubled by feelings of guilt?’. The responses ‘I don’t know’ and ‘I do not wish to answer’ were recoded to missing data because they do not provide any information on the latent trait of neuroticism.

### Analytic strategy

An item-response theory (IRT) model was used to investigate item characteristics. Details of the IRT model are specified in the supporting information. In brief, the IRT model describes how items contribute to the assessment of a latent trait, such as neuroticism. This parameter is by convention called θ and is standardised to a mean value of zero and a range of −4 to +4 standard deviation units. For each item, a difficulty parameter (*α*) identifies which level of θ the item most efficiently describes. For example, in the EPQ-RN which items are most likely to be endorsed as “Yes” by individuals with high neuroticism.

Also, for each item, a discrimination parameter (γ) describes how well each item discriminates between different levels of θ. For example, in the EPQ-RN, is an item scored “Yes” only by those with high neuroticism, or also by those with moderate levels of neuroticism. Difficulty is measured as the point of inflection of a logistic regression curve between “Yes” and “No scores where high scores reflect greater difficulty. Discrimination is estimated as the slope of the inflection point between “Yes” and “No scores, where higher values of β reflect greater discrimination. Difficulty and discrimination parameters are used to select items that collectively have high levels of discrimination across a range of θ values, rather than clustering around a single value [14].

The scalability of items, i.e. the extent to which they provide a unidimensional monotonic scale was assessed by Mokken analysis, where high values of Loevinger’s H between 0.5 and 1 suggest high scalability.

To explore the potential for improving the psychometric properties of the EPQ-RN, a backwards stepwise approach to item removal was adopted. The goal was to identify items covering a broad range of difficulty, with high levels of discrimination, and high scalability scores.

UK Biobank data for this analysis (application 15697) were uploaded onto the Dementias Platform UK (DPUK) Data Portal [15] and analysed using STATA 17.0 [16].

## Results

### Sample

The entire UK Biobank sample available after withdrawals and with complete EPQ-RN data was included in the analysis (*n* = 401,527). Participants were aged 39 to 73 years (*M* = 56.41 years; *SD* = 8.07, 53.68% female).

### IRT analysis

For the 12-item scale, difficulty ranged between α = −0.14 and α = 1.41. with 6 items clustering between 0 and 1 (Table 1). These findings can be visualised using item characteristic curves and item information functions (Fig 1). Both plots show the EPQ-RN to be moderately efficient for measuring the middle range of neuroticism, and that high scores are a cumulation of middle range item scores, rather than items which are sensitive to high (or low) levels of neuroticism. This suggests a degree of measurement duplication.

**Table 1.** IRT model item parameters for the 12-item scale.

| Item | Parameter |  |  |
| --- | --- | --- | --- |
| Mood go up and down? | .21 | .28 | .39 |
| Feelings easily hurt? | -0.13 | .60 | .39 |
| Are you a worrier? | -0.13 | .85 | .44 |
| Suffer from nerves? | .17 | .67 | .44 |
| Feel miserable for no reason? | .29 | .97 | .35 |
| Often feel fed-up? | .36 | .09 | .39 |
| Tense or highly strung? | .23 | .05 | .43 |
| Often feel lonely? | .41 | .47 | .46 |
| An irritable person? | .95 | .34 | .41 |
| A nervous person? | .03 | .85 | .43 |
| Worry embarrassing experience? | .15 | .45 | .47 |
| Troubled feelings of guilt? | .86 | .54 | .47 |

Notes: Item names truncated for brevity, see text. $\alpha$ Item difficulty, $\beta$ Item discrimination, H
Loevinger Coefficient, $p < .0001$ for all $\beta$ values; Goodness of fit (root mean square error of
approximation) = 0.11

**Fig 1.**
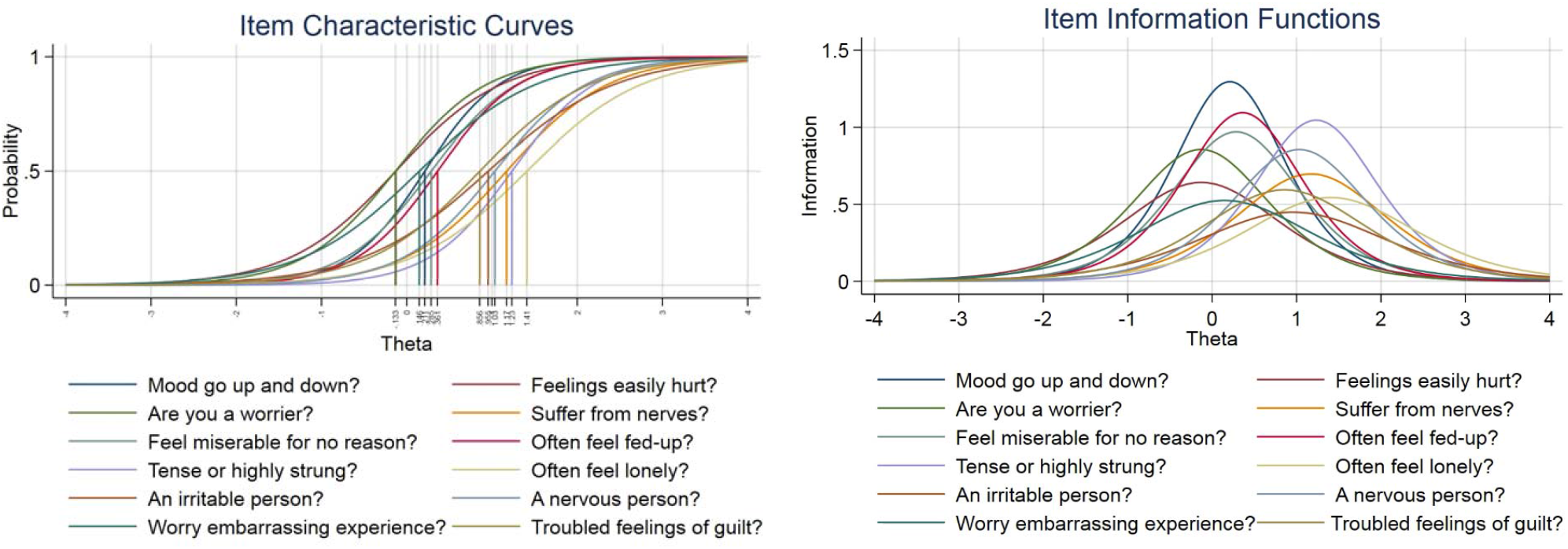
Item Characteristic Curves (ICC) and Item Information Function (IIF) graph for the 12-item scale.

Discrimination parameters ranged between β = 1.34 and β = 2.28. The item measuring ‘Does your mood often go up and down?’ exhibits the highest level of discrimination at 2.28, suggesting that this ‘mood’ question possesses the highest amount of information synonymous with the neurotic trait. In contrast, the item ‘Are you an irritable person?’, 1.34, is the lowest, and below the recommended level of 1.7 for measuring trait values [14]. The items, ‘Are you a worrier?’; ‘Do you ever feel just miserable for no reason?’; ‘Do you often feel fed-up?’ and ‘Would you call yourself tense or highly strung’ and ‘Would you call yourself a nervous person?’ had discrimination values of above 1.7.

For scalability, the monotonicity parameter, Loevinger’s H, ranged between 0.35 and 0.47 with four items scoring < 0.4, indicating poor scalability. This combination of a limited α range, modest β scores, and relatively low H values describes a relatively inefficient instrument. This is confirmed by poor goodness of fit (RMSEA = 0.11).

The potential for improving the psychometric properties of the EPQ-RN was limited by the relatively narrow range of item difficulty scores; there being no items particularly sensitive to high or low levels of neuroticism (Fig 1). This constrained our item selection strategy to identifying least discriminating items, and omitting items with identical difficulty scores according to scalability and goodness of fit. In order of removal, omited items were ‘Are you an irritable person?’ (β = 1.34), ‘Do you often feel lonely?’ (β = 1.49), ‘Are you often troubled by feelings of guilt?’ (β = 1.54), ‘Do you worry too long after an embarrassing experience?’ (β = 1.45), and ‘Are your feelings easily hurt?’ (β = 1.43). For the remaining seven items the discrimination scores ranged between 1.57 and 2.57, the difficulty score ranged between α = −0.15 and α = 1.25, with all items showing moderate to good scalability with H values 0.47 to 0.51 (Table 2). However, the 7-item scale did not show an improved goodness of fit (RMSEA = 0.17). ICC and IIF plots for the 7-item scale can be found in S1 Fig.

To explore further efficiencies the item elimination process was continued until there were 3 remaining items, each with non-duplicative difficulty scores, high discrimination scores and high scalability scores (Table 2). Goodness of fit for the 3-item scale was high (RMSEA = 0.00). ICC and IIF plots for the 3-item scale can be found in S2 Fig.

**Table 2.** IRT model item parameters for the 7-item and 3-item scales.

| Item | 7-item scale parameters |  |  | 3-item scale parameters |  |  |
| --- | --- | --- | --- | --- | --- | --- |
| | $\alpha$ | $\beta$ | H | $\alpha$ | $\beta$ | H |
| **Mood go up and down?** | 0.20 | **2.57** | 0.50 | 0.19 | 3.40 | 0.57 |
| **Are you a worrier?** | **-0.15** | **1.57** | 0.51 | - | - | - |
| **Suffer from nerves?** | 1.16 | 1.70 | 0.47 | - | - | - |
| **Feel miserable for no reason?** | 0.27 | 2.16 | 0.47 | 0.26 | 2.77 | 0.54 |
| **Often feel fed-up?** | 0.35 | 2.20 | 0.47 | 0.33 | 2.89 | 0.56 |
| **Tense or highly strung?** | **1.25** | 1.98 | 0.52 | - | - | - |
| **A nervous person?** | 1.05 | 1.80 | 0.49 | - | - | - |

*Notes:* Item names truncated for brevity, see text. $\beta$ Item discrimination, $\alpha$ Item difficulty, H Loevinger Coefficient $p < .0001$ for all $\beta$ values; 7-item Goodness of fit (root mean square error of approximation) = 0.17; 3-item Goodness of fit (root mean square error of approximation) < 0.001.

An important issue is the comparability of assessment between the 12, 7 and 3 item scales. To assess this estimated θ scores for each individual were calculated and ranked for each scale. The intraclass correlation rank correlation between the 12-item and 7 item scale was r = 0.95, indicating 90.25% agreement between scales. Between the 12-item and 3-item scales the correlation was r = 0.84, whilst between the 7 item and 3-item scales r = 0.91.

The reliability of the scale and statistical assumptions of the IRT models are reported in the supporting information.

In summary, the overall pattern of item distribution across the θ continuum suggests that across the 12-item EPQ-RN neuroticism scale there are no items which measure an extreme level of neurotic trait characteristics or an extreme level of non-neurotic trait characteristics. It suggests that the questions are mostly measuring the neurotic trait characteristics which have a higher probability of endorsement by individuals who are experiencing a minimal to no level of neuroticism (θ = −0.13 to 1.41).

## Discussion

In a large population cohort of 401,527 adults aged 39-73 years, limitations in the range and reliability of item trait characteristics were found across the 12-item EPQ-RN scale when an IRT model was estimated. Our findings suggest that the 12-item scale is inefficient with poor discrimination and scalability at the extreme ends of the scale range, such that high and low trait levels are poorly assessed. A reliability function analysis suggests there is poor reliability at the extremes of the scale score and high neuroticism scores derived from the EPQ-RN are a function of accumulative mid-range values. Through systematic item deletion and mathematical assessment, a revised 7-item version of the scale with greater item discrimination and reliability was found, suggesting that selective items within the 12-item version are redundant. A further reduced 3-item version was investigated but although this scale possesses items of high discrimination and scalability, item range is very narrow (α = 0.19 - 0.33) and lacks reliability.

To our knowledge, this is the first study to conduct a comprehensive psychometric scale assessment applying IRT to the EPQ-RN on such a large population. IRT has been successfully used to assess the item efficiency in psychiatric scales such as the 16-item Anxiety Sensitivity Index [17] and the 10-item feelings scale for depression [18] and it is increasingly being adopted to revise existing healthcare scales, such as the Simple Clinical Colitis Activity Index [19]. The reduction and choice of items however, is not a clearly defined process, notwithstanding the emergence of criteria for mathematical assumptions such as the 1.7 discrimination guidance [14] and Loevinger H criteria for scalability [20]. These criteria are simultaneously taken into account with the estimated IRT model output, theoretical understanding of the construct of interest and scale application. For example, it might not be beneficial to have a short 3-item scale if all items are highly discriminatory around the same value of θ and no information is provided about patients or participants who lie along the rest of the θ scale (−4 to +4). Reducing patient and participant burden needs to be weighed up against item reduction and, scale design and purpose.

Utilising psychometric methodologies to analyse psychosocial and health-related outcomes has important implications for analysing longitudinal change both in clinical settings and epidemiological research. An IRT analysis provides item-level information and scaling characteristics through the further computation of post-estimation assumptions including the estimation of an individual θ latent metric predictive of individual θ scores on the fitted IRT model. This θ metric may then be used as a latent construct in assessing longitudinal change [21] which may be a more reliable measure compared to a single summated score [22]. Furthermore, it has also been suggested that using an IRT derived θ in longitudinal studies, over the summated score, may be preferable with reducing overestimation of the repeated measure variance and underestimation of the between-person variance [23].

A further advantage of utilising psychometric methodologies in an epidemiological context is that IRT extends the opportunity to utilise, computer adaptive testing (CAT) for both scale development and for efficient test delivery. During CAT administration, θ is automatically computed in response to the trait (θ) of the respondent and it is therefore not necessary to present the full range of items as the response scale is adaptive to individual performance (trait level), the items underlying the trait and a stopping rule [24]. The potential to reduce a scale so that only the most reliable and informative questions are presented to participants is essential in clinical settings and epidemiological research. This is important to consider when working with individuals who are older or who have co-comorbid psychiatric disorders. Moreover, focused, reliable and user-friendly scales in a research setting increase user satisfaction, reduce participant burden and maintain long-term participant retention.

Participants who display or possess the extreme trait characteristics are rare, however, the potential should exist for this eventuality, but many scales are simply not adequately designed to do so [21]. Moreover, previous research suggests that both the 12 and 3-item EPQ-RN neuroticism scales may have reduced power to discriminate between low and high scoring individuals [12]; we found evidence of this in the 12-item scale. It is important in both clinical and research settings that scales are designed to measure across the trait spectrum and this is possible if scales are developed using psychometric methodologies such as those described here and elsewhere [25, 26]. Future research and scale development should, therefore, develop a neuroticism scale that measures the entire latent trait continuum (θ) by also including items with high difficulty, i.e. items that only individuals with a high latent trait would endorse, and by also including items measuring the opposites of trait neuroticism, such as emotional stability. Further research is also needed to validate the 7-item EPQ-RN scale and to investigate its construct validity by comparing the scale to other establishes measures of neuroticism such as the NEO five-factor personality inventory neuroticism scale [27].

## Conclusions

The 12-item neuroticism EPQ-R scale lacks item reliability and neurotic trait-specific information at the extreme ends of the neurotic continuum when an IRT model is estimated. A secondary analysis suggests that systematic item-elimination and re-estimation of the model produces a 7-item scale with higher levels of item information and reliability. This study suggests that the 12-item EPQ-R scale could benefit from item revisions and updating including item deletions. Strengths of this study were the large population cohort available for a comprehensive IRT analysis and the psychometric methodologies which were applied to the data.

## Acknowledgements

All analyses were conducted on the Dementias Platform (DPUK) Data Portal using UK Biobank application 15697 PI John Gallacher for DPUK project 0169. The Medical Research Council supports DPUK through grant MR/T0333771. Sarah Bauermeister and Patrick Pflanz are supported by DPUK. Access to the data can be requested through UK Biobank (https://www.ukbiobank.ac.uk/enable-your-research/apply-for-access).

## Declarations

### Financial Disclosure Statement

The Medical Research Council supports DPUK through grant MR/T0333771

### Competing interests

SB, CPP and JG declare no competing interests

### Ethics approval and consent to participate

Analysis of secondary data only with ethical approval in place from source cohort, UK Biobank Research Ethics Committee - REC reference 11/NW/0382.

### Consent for publication

SB, CPP and JG give full consent for publication

### Availability of data and materials

The dataset(s) supporting the conclusions of this article is(are) available in the Dementias Platform UK (DPUK) Data Portal repository, https://portal.dementiasplatform.uk/.

### Authors’ contributions

SB and JG conceptualised the idea. CPP and SB analysed and interpreted the data, and wrote the manuscript. CPP and JG edited and proofread the manuscript. All authors read and approved the final manuscript.

## Supporting information

### Supporting methods

For these binary response data, a 2-parameter logistic (2-PL) IRT model is appropriate:

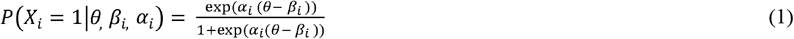

The dependent variable is the dichotomous response (yes/no), the independent variables are the person’s trait level, theta (θ) and item difficulty (*β_i_*). The independent variables combine accumulatively and the item’s difficulty is subtracted from θ. That is, the ratio of the probability of success for a person on an item to the probability of failure, where a logistic function provides the probability that endorsing any item (*i*) is independent from the outcome of any other item, controlling for person parameters (θ), and item parameters. The 2-PL model includes two parameters to represent the item properties (difficulty and discrimination) in the exponential form of the logistic model. Previous research showed that a 2-PL IRT model was appropriate for Eysenck scales [28].

For each item, an item response function (IRF) may be calculated which calibrates the responses of an individual against each item. A calibrated standardised score for trait severity θ is returned and may be plotted as item characteristic curves (ICC) along a standardised scale with a mean of 0 (see Fig 1). From the ICC two parameters may be estimated. The first is the value of θ at which the likelihood of item endorsement is 0.5, interpreted as ‘expressed trait severity’. The second is the slope of the curve from the point at which the likelihood of item endorsement is 0.5, interpreted as ‘expressed item discrimination’ i.e., the ability to discriminate between greater and lesser severity scores. The IRF may also be expressed as an item information curve (IIF) which displays the relationship between severity and discrimination (see Fig 1). The apex of the curve for any IIC indicates the value of θ at which there is maximum discrimination. Statistical assumptions underlying the IRT principles of scalability, unidimensionality and item independence are examined.

### Supporting results

#### Statistical assumptions of the IRT analysis for the 12-item scale

##### 1. Item independence

A correlation analysis assessed initial item independence and all items were significantly correlated (*p* < .0001) but the majority of values were lower than 0.50, suggesting basic local item independence. A residual coefficient matrix, requested after estimation of a single-factor model showed that no residuals were too highly correlated, *R* < 0.20 [29], suggesting basic item independence.

##### 2. Monotonicity

A Mokken analysis produced a Loevinger H coefficient [30] which measures the scalable quality of items, expressed as a probability measure, independent of a respondent’s θ. These coefficients ranged between 0.35 and 0.47, suggesting a weak (H = 0.3-0.4) to moderate (H = 0.4-0.5) monotonicity, no items reached strong scalability (H ≥ 0.5) [30].

##### 3. Unidimensionality

A single-factor CFA model was used to test for unidimensionality of the 12-item EPQ-N scale. The single-factor model had poor model fit: Chi^2^ = 241797.89, p < 0.0001, Root Mean Square Error of approximation (RMSEA) = 0.11, Comparative Fit Index (CFI) = 0.81, Tucker-Lewis Index (TLI) = 0.76, thereby indicating that the scale did not fulfill strict criteria for unidimensionality. Since IRT is relatively robust to violations against the assumption of unidimensionality [31, 32], the violation of unidimensionality is not a major concern when estimating item characteristics from IRT in the following. A post-IRT estimation model measure of unidimensionality was also computed using a semi-partial correlation controlling for θ. This analysis provides individual item variance contribution after adjusting for all the other variables including θ. It demonstrates the relationship between local independence and unidimensionality, reflecting a conservative assessment whereby the desired *R*^2^ should ideally be zero or as close to zero as possible [33]. Items ranged between 0.01 and 0.02, suggesting unidimensionality. To our knowledge, there is still no standardised cut-off criterium for assessing this value (i.e., how close to zero all items should be across a scale).

#### Statistical assumptions of the IRT analysis for the 7-item scale

Statistical assumptions were computed on the revised scale of 7 items and importantly a Mokken analysis suggests improved scalability (monotonicity) compared to the full 12-item scale with three items reaching values ≥ 0.50. Acceptable metrics for unidimensionality and item independence were achieved for this revised scale. A single-factor CFA model showed that the 7-item scale did not fulfill strict criteria for unidimensionality: Chi^2^ = 153492.42, p < 0.0001, RMSEA = 0.17, CFI = 0.79, TLI = 0.68.

#### Statistical assumptions of the IRT analysis for the 3-item scale

A Mokken analysis suggests that scalability is strong (H ≥ 0.50) across all items. A single-factor CFA model showed that the 3-item scale was unidimensional: Chi2 = 0, p = 1, RMSEA = 0, CFI = 1, TLI = 1. In a semi-partial correlation analysis controlling for θ, item R^2^ ranged between 0.09 and 0.13, suggesting basic local independence and unidimensionality.

#### Reliability of the scales

##### Reliability of the 12-item scale

In IRT, reliability may be calculated at multiple point values of θ along the continuum rather than a single reliability score as in CTT. Reliability is defined at different points of θ with the mean of θ fixed at 0 and the variance at 1, facilitating identification of the model and reliability for all points along the θ continuum, distinguishing respondents according to specific values of θ [29]. For the 12-item scale, there is reliable information to differentiate respondents who possess no or just above an average amount of trait information (θ=0; 0.87 and θ=1; 0.88), considered very good for reliability However, reliability then decreases (θ=2; 0.76 and θ=−1; 0.71) suggesting that the highest reliability of measuring the neurotic trait is at normal or a minimal amount of neuroticism, θ=0 or 1. Thereafter, reliability reduces so that the extreme end of the continuum, θ=3; 4; −2; −3; −4, is no longer reliably measured (S1 Table).

S1 Table. Reliability for values of θ from the 2-PL IRT model fit for the 12-item, 7-item and 3-item scales.

##### Reliability of the revised scales

Reliability across the revised 7-item scale is marginally improved compared to the full scale suggesting redundancy of the removed items (S1 Table). Reliability across the revised 3-item is only good at θ = 0 suggesting this scale is only reliable to measure those with an average trait (S1 Table).

### Supporting tables

**S1 Table.** Reliability for values of θ from the 2-PL IRT model fit for the 12-item, 7-item and 3-item scales.

| $\Theta$ | 12-item scale | | | 7-item scale | | | 3-item scale | | |
| --- | --- | --- | --- | --- | --- | --- | --- | --- | --- |
|  | TIF | TIF SE | Reliability | TIF | TIF SE | Reliability | TIF | TIF SE | Reliability |
| **-4** | 1.02 | 0.99 | 0.02 | 1.01 | 1.00 | 0.01 | 1.00 | 1.00 | 0.00 |
| **-3** | 1.10 | 0.95 | 0.09 | 1.04 | 0.98 | 0.04 | 1.00 | 1.00 | 0.00 |
| **-2** | 1.52 | 0.81 | 0.34 | 1.24 | 0.90 | 0.19 | 1.03 | 0.98 | 0.03 |
| **-1** | 3.43 | 0.54 | 0.71 | 2.36 | 0.65 | 0.58 | 1.59 | 0.79 | 0.37 |
| **0** | 7.96 | 0.35 | 0.87 | 6.22 | 0.40 | 0.84 | 6.96 | 0.38 | 0.86 |
| **1** | 8.04 | 0.35 | 0.88 | 5.82 | 0.41 | 0.83 | 3.34 | 0.55 | 0.70 |
| **2** | 4.11 | 0.49 | 0.76 | 2.84 | 0.59 | 0.65 | 1.15 | 0.93 | 0.13 |
| **3** | 1.77 | 0.75 | 0.44 | 1.37 | 0.85 | 0.27 | 1.01 | 1.00 | 0.01 |
| **4** | 1.16 | 0.93 | 0.14 | 1.06 | 0.97 | 0.06 | 1.00 | 1.00 | 0.00 |

*Note:* TIF = Test Information Function; SE = standard error.

### Supporting figures

**S1 Fig.**
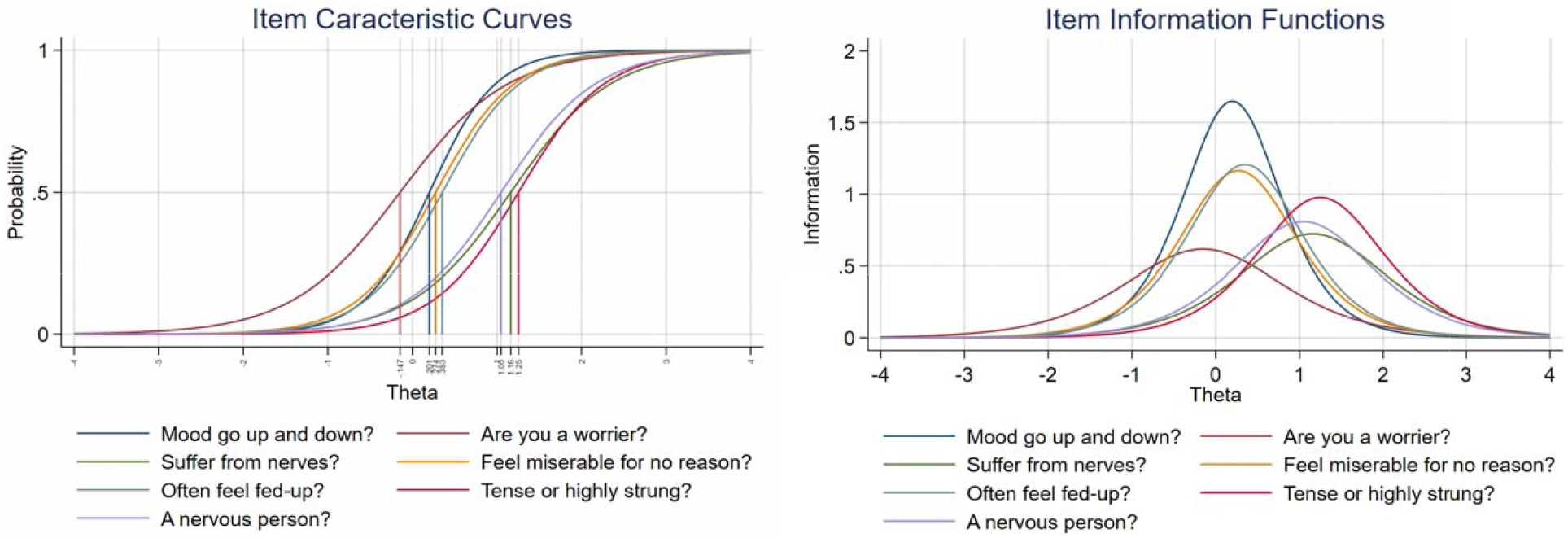
Item Characteristic Curve (ICC) and Item Information Function (IIF) graph for the 7-item scale.

**S2 Fig.**
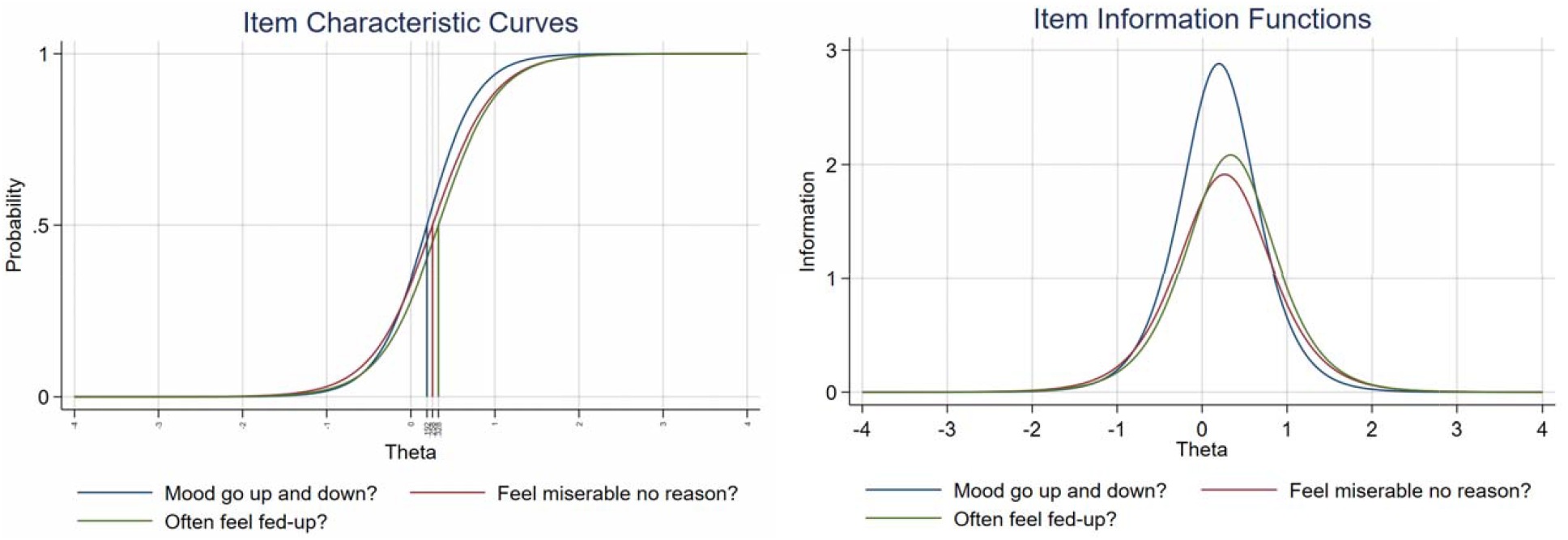
Item Characteristic Curves (ICC) and Item Information Function (IIF) graph for the 3-item scale.

## Notes

### Competing Interest Statement

The authors have declared no competing interest.

### Summary of Updates

The data released by UK Biobank has been renewed with current withdrawals. The manuscript has been revised to be more succinct but the overall message of the manuscript is the same.

